# Molecular characters and phylogenetic analysis of *Clostridium perfringens* from different regions in China, from 2013 to 2021

**DOI:** 10.1101/2022.09.23.509295

**Authors:** Jia xin Zhong, Hao ran Zheng, Yuan yuan Wang, Lu lu Bai, Xiao li Du, Yuan Wu, Jin xing Lu

## Abstract

*Clostridium perfringens* (*C. perfringens*) is a significant foodborne pathogen and a common cause of intestinal diseases in both animals and humans. Altogether, 186 isolates were obtained from humans (n = 147), animals (n = 25), and food (n = 14), comprising 174 type A strains (93.55%), 11 type F strains (5.91%), and one type D strain (0.54%); and were analyzed by multilocus sequence typing (MLST) and antimicrobial susceptibility testing. Additionally, some specific ST complexes were analyzed by cgMLST and cgSNP to investigate genetic relatedness. MLST indicated the most prevalent STs of *C. perfringens* of human and animal origin were as follows: ST221 (5/147), ST62 (4/147), ST408 (4/147), and ST493 (4/147) were predominant in humans, while ST479 (5/25) was the major type in animals. Within the same ST complex, genetically unrelated relationships or potential clustering/transmission events were further recognized by cgMLST and cgSNP, illustrating that these two methods are valuable in defining outbreaks and transmission events. All tested isolates were susceptible to vancomycin and meropenem. The rates of resistance to metronidazole, penicillin, cefoxitin, moxifloxacin, and clindamycin were low (metronidazole: 1.08%; penicillin: 9.68%; cefoxitin: 0.54%; moxifloxacin: 6.45%; and chloramphenicol: 3.76%). Interestingly, 49.66% of human origin were clindamycin-resistant, and 18.2% were penicillin-insensitive. Importantly, the portion of MDR isolates was significantly lower than in previous reports. The study provides an overview of the epidemiological characteristics of *C. perfringens* with different origins and hosts in China. *C. perfringens* demonstrated remarkable genetic diversity and distinct molecular features compared to antibiotic-resistance profiles from other studies.

**IMPORTANCE:** *C. perfringens* is one of the most common bacterial causes of foodborne illness globally, responsible for several food-poisoning outbreaks each year. This study provides an overview of *C. perfringens* isolates from different hosts and regions in China according to MLST, antibiotic-resistance characters, cgMLST, and cgSNP analyses, showing high genetic diversity and identifying potential clustering and transmission events. The antimicrobial profile in this study was distinct from that of a previous report with a much lower MDR rate, indicating that *C. perfringens* in China needs further investigation.

## Introduction

*Clostridium perfringens* (*C. perfringens*) is a Gram-positive anaerobic bacterium that is abundant in nature. The bacteria can cause gas gangrene, intestinal diseases, and food poisoning in both animals and humans (1, 2). The significant pathogenic mechanism of *C. perfringens* is the production of a wide variety of toxins and enzymes (3). *C. perfringens* is classified into seven types: A (ɑ), B (ɑ, b, ƹ), C (ɑ, b), D (ɑ, ƹ), E (ɑ, ι), F (ɑ, cpe), and G (ɑ, netB) by a toxin-based typing scheme (4).

Antibiotic-associated diarrhea (AAD) continues to be a major healthcare issue, both in hospitalized patients and in the community (5). Many investigations have revealed that *C. perfringens* played an important role in diarrheic patients with a history of antibiotic use (6, 7). According to some studies, *C. perfringens* is responsible for 5 to 20% of all cases of AAD and sporadic non-foodborne diarrhea (6, 8). Recently, an increase in *C. perfringens* antimicrobial resistance has been noted, and some investigations have indicated that most *C. perfringens* strains of animal origin are multidrug resistant (MDR) (9, 10). However, in China the research on *C. perfringens* antimicrobial resistance is limited.

The technique of MLST is widely applied for the identification of human, animal, and foodborne pathogens, and it has also been used to analyze bacterial population diversity and investigate *C. perfringens* outbreaks (11). With the increasing application of whole-genome sequencing (WGS), high-resolution molecular subtyping approaches such as core genome MLST (cgMLST) based on WGS have gained popularity in epidemic investigation and bacterial tracing (12, 13). Previous research has shown that *C. perfringens* from different sources had a large amount of genetic variation (14-16), and human strains demonstrated more variation than animal strains (17). However, the molecular characters of *C. perfringens* isolates from different sample origins and regions in China are limited. Therefore, this research aimed to provide valuable epidemiological data on genotype distribution, antibiotic resistance, toxin type, genetic diversity, and phylogenetic characteristics of *C. perfringens* from distinct geographical regions in China. The findings can provide a deeper understanding of the molecular epidemiology of *C. perfringens* in China and offer a basis for the development of rapid detection and surveillance networks.

## Results

### Evolution and phylogenetic analysis of *C. perfringens* isolates in China by MLST

MLST analysis revealed that 186 *C. perfringens* isolates exhibited 135 sequence types (STs) with a high degree of genetic diversity, including 93 new STs. In the segments of *colA* (18), *groEL* (16), *sodA* (20), *plc* (23), *gyrB* (13), *sigK* (10), *pgk* (9), and *nadA* (20), up to 129 new alleles were identified, resulting in new genotypes. Among the 132 STs, the most frequent ST was ST221 (7/186, 3.74%), followed by ST408 (5/186, 2.67%), ST479 (5/186, 2.67%), ST62 (4/186, 2.14%), and ST493 (4/186, 2.14%). There were 98 STs containing only one isolate. The 186 strains were from humans (n = 147), animals (n = 25) and food (n = 14) (Fig. 1). The human strains comprised 110 STs, with ST221 (5/147), ST62 (4/147), ST408 (4/147) and ST493 (4/147) being the predominant STs. The animal strains contained 19 STs, with ST 479 (5/25) and ST476 (2/25) being the predominant STs (Fig. 2). Supplementary Table S1 contains a complete list of STs found in this study.

**Figure 1.**
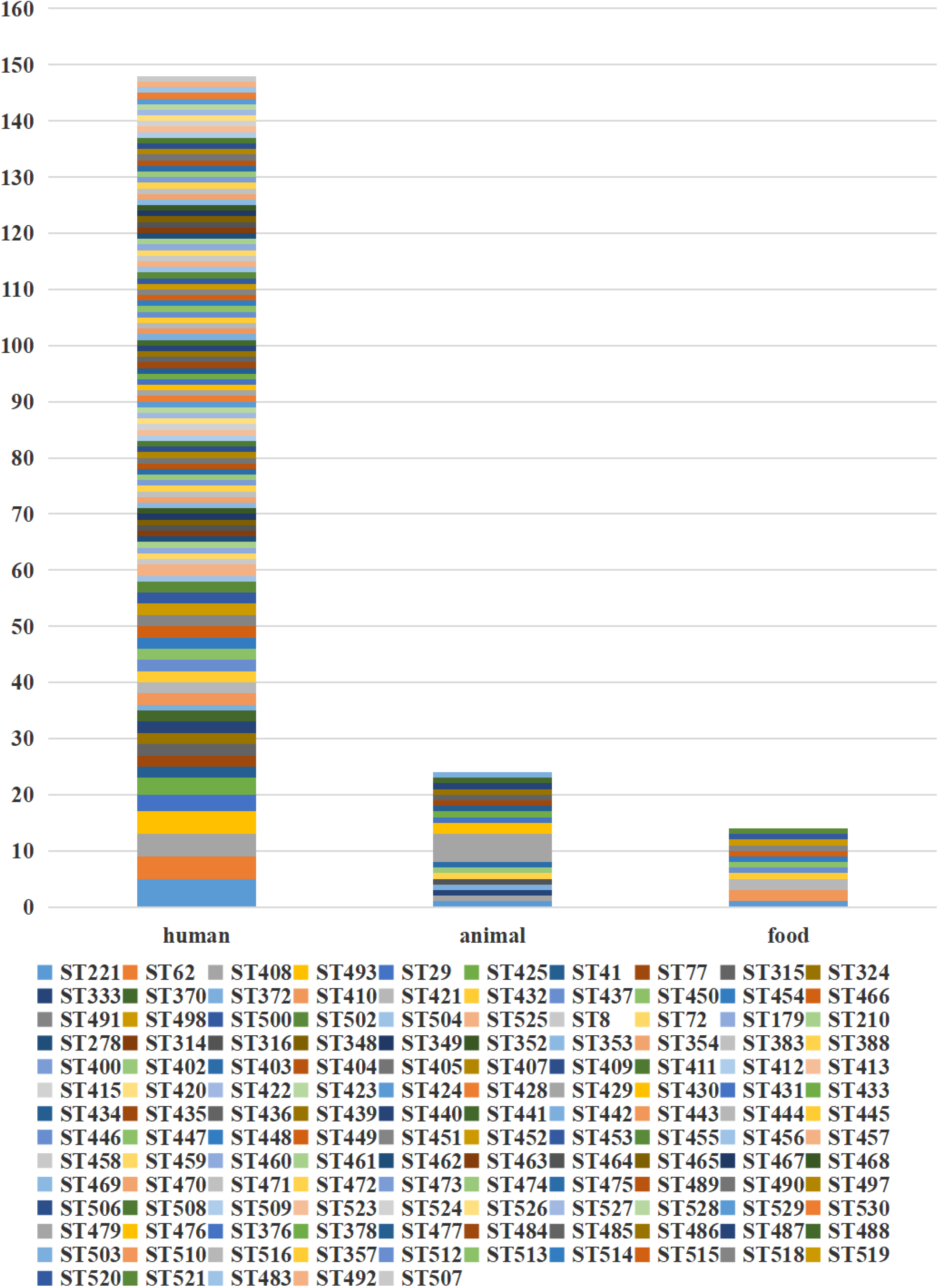
Isolation distribution of 186 *C. perfringens* isolates.

**Figure 2.**
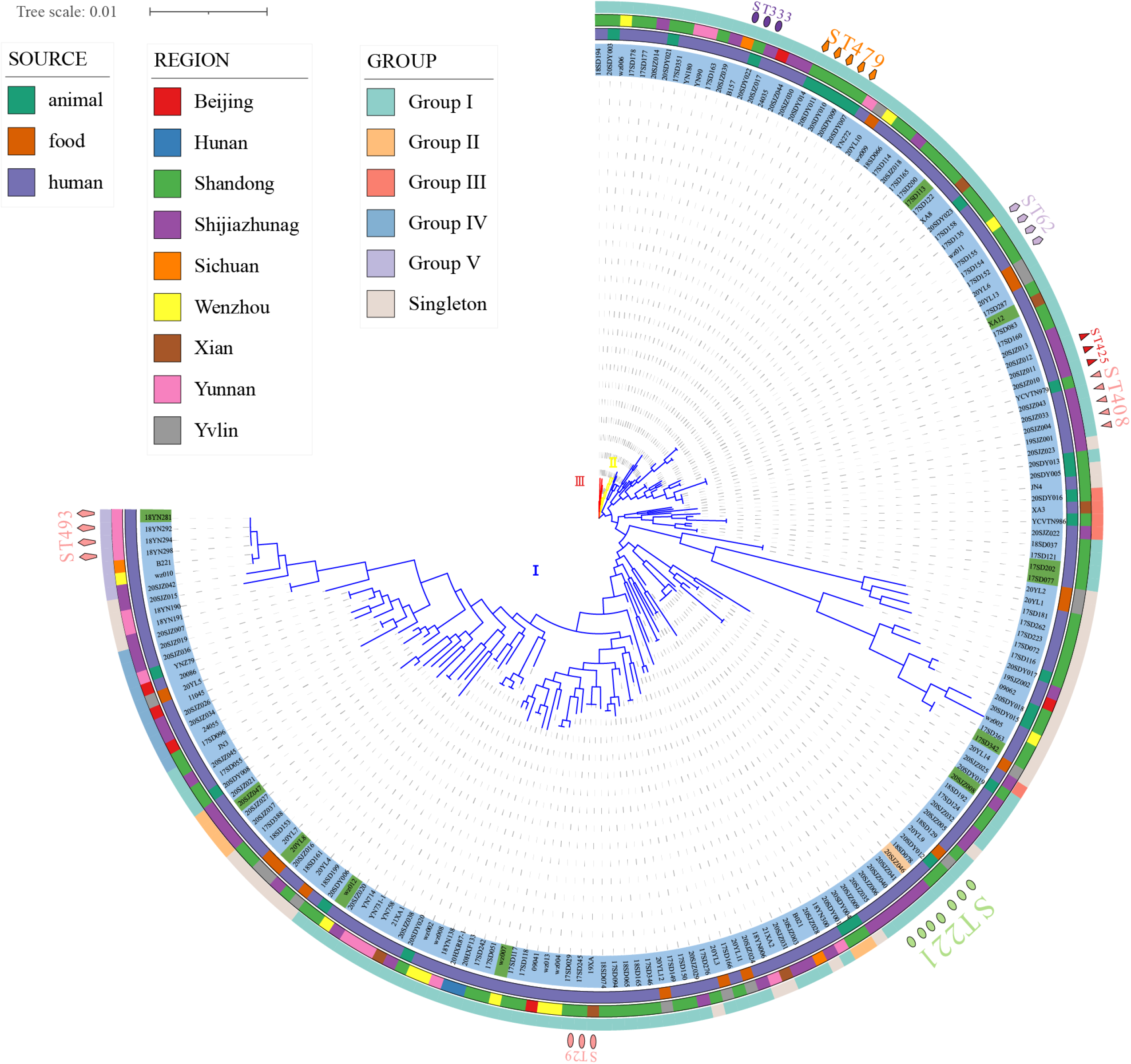
Phylogenetic maximum likelihood tree based on MLST identified in 186 strains of *Clostridium perfringens* (bootstrap test = 1000 replicates) by fasttree. The figure shows the population structure of the strains with toxin type next to the ML tree (blue, type A; green, type F; and orange, type D), followed by the region, source, and ST of isolation as in the legend. The phylogenetic tree was visualized using iTOL.

Of the 186 total examined isolates, 174 (93.55%) belonged to type A; 11 isolates (5.91%) were classified to type F; one isolate (0.54%) was classified to type D (Fig. 2); and none of the investigated isolates were types B, C, and/or E. Clustering of the 186 genomes revealed that 91.94% (n = 171) belonged to phylogroup I, followed by phylogroups II (n = 10) and III (n = 5) (Fig. 2). Phylogroup I, the most abundant subgroup, contained all predominant STs (ST221, ST408, ST479, ST62, and ST493), all toxin types (A, F, D), and all sources and all regions (Fig. 2). As described, phylogroup I was associated with isolates from humans, food, and animals; and this group had a broad ecological distribution (Fig. 2). The strains of toxin type F and toxin type D were also concentrated in phylogroup I (Fig. 2). Phylogroup II contained ST333 (n = 3) as the predominant type and also included ST372 (n = 2) and five unique STs (Fig. 2). Phylogroup III contained five isolates with different STs (Fig. 2). Two animal isolates and eight human isolates were included in phylogroup II, while one animal isolate and four human isolates were included in phylogroup III (Fig. 2). Strains in phylogroup II and phylogroup III were all type A.

There were no significant host, geographic distribution, or toxin type connections among the strains involved (Fig. 2). In the phylogenetic maximum likelihood tree, there were several STs, including isolates with different regions, hosts, or different toxin types within the same ST such as ST221, ST408, ST333, ST372, ST316, ST388, ST402, and ST475, indicating that isolates may have potentially spread across different hosts or regions (Fig. 2). For example, ST221 contained seven strains from clinical diarrhea patients [Shijiazhuang (n = 4), Shandong (n = 1)], sheep [Shandong (n = 1)] and food [Yulin (n = 1)] (Fig. 2). ST408 consisted of five strains originating from AAD patients [Shijiazhuang (n = 4)] and a calf [Shandong (n = 1)] (Fig. 2).

A total of five groups (Group I–Group V) were found in the minimal spanning tree (Fig. 3) among the studied strains isolated in this investigation, accounting for 82.80% (154/186) of the strains. Thirty-two STs were determined to be singletons, with no observed cluster relationships. Group I, the most prolific cluster, included 80 STs (including advantage STs ST221, ST408, ST479, ST62, ST333, and ST493), with a total of 122 strains that accounted for 65.59% (122/186) of all detected strains. Group II contained four clinical origin strains (ST412, ST413, ST439, and ST441) and two animal origin strains (ST476), accounting for 3.23% (6/186) of the examined strains. Group III contained three clinical origin strains (ST402, ST431, and ST524) and two animal origin strains (ST402 and ST485), accounting for 2.69% (5/186) of the examined strains. Group IV contained eight human origin strains (ST410, ST432, ST447, ST526, ST527, and ST529), one animal origin strain (ST503), and one food origin strain (ST514), accounting for 5.38% (10/186) of the examined strains. Group V contained seven human origin strains (ST440, ST493, ST504, and ST507), accounting for 3.76% (7/186) of the examined strains; these showed significant genetic distance (≥ 6 allele differences) with other isolates in the minimum-spanning tree (Fig. 3) and significant genetic distance (≥7 allele differences) with other groups (Fig. 3). Half of the food origin strains were dispersed as singletons, while most strains with animal origin clustered with strains of human origin, indicating the character of the zoonotic food-borne pathogen *C. perfringens*.

**Figure 3.**
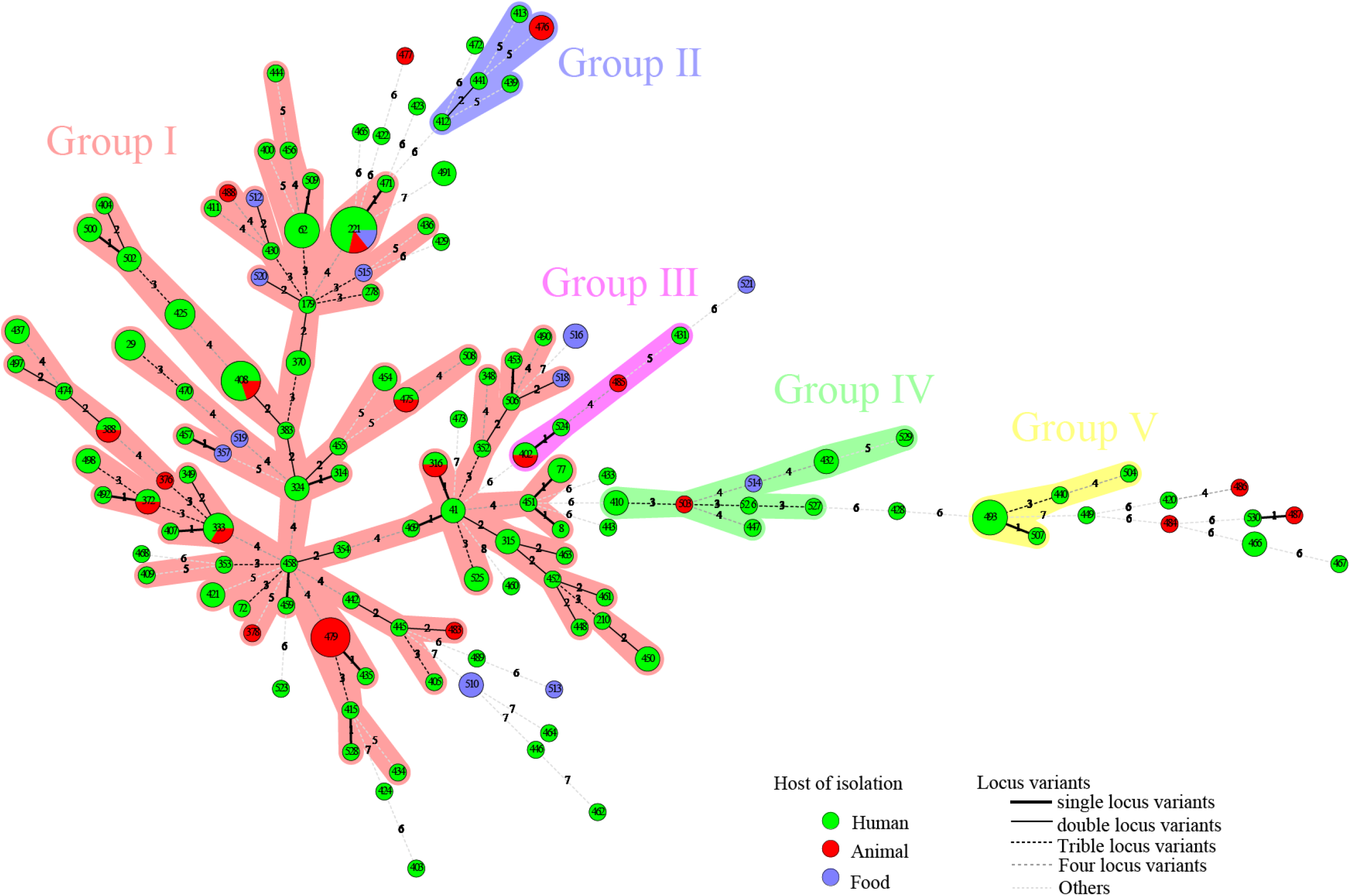
A total of 186 strains of *C. perfringens* from different sources were analyzed by constructing an MLST-minimal spanning tree. The minimum-spanning tree was constructed using the Bionumerics software (Bionumerics, version 5.1). The shaded section represents five clusters. The clusters were assigned for at least three types within five changes of neighbor distance. The area of the circle represents the number of strains; different colors represent different hosts; and the number on the branch represents the difference compared to housekeeping genes.

Interestingly, the distribution of isolates from multiple sources in the minimum-spanning tree (Fig. 3) and in the phylogenetic tree (Fig. 2) was not consistent. Due to different mathematical calculation models (the minimum-spanning tree was based on a matrix, while the phylogenetic tree was based on nucleic acid sequences), some strains belonging to similar branches in the phylogenetic tree did not form the same clusters in the minimum-spanning tree. For example, some isolates in phylogroup I were assigned to different clusters of the minimum-spanning tree (Figs. 2 and 3).

### cgMLST and cgSNP evolutionary relationship analysis of the same STs

Figure 3 shows the genetic relatedness of strains in the same STs on the cgMLST scheme from PubMLST based on the presence/absence of 1,431 highly conserved core genes.

Isolates with the same ST221 were divided into seven cgMLST types, comprising five from clinical human isolates, one from food, and one from animals (Fig. 4A). The seven genomes showed ≤ 100 allele differences (range 20–100). Furthermore, the food isolate cp2020YL9 (ST221) and sheep isolate cp2020SDY012 (ST221) showed 35 and 100 allele differences, respectively, compared to the human cp2018SD078 (ST221) isolate, indicating that these isolates with identical ST were genetically unrelated (Fig. 4A).

**Figure 4.**
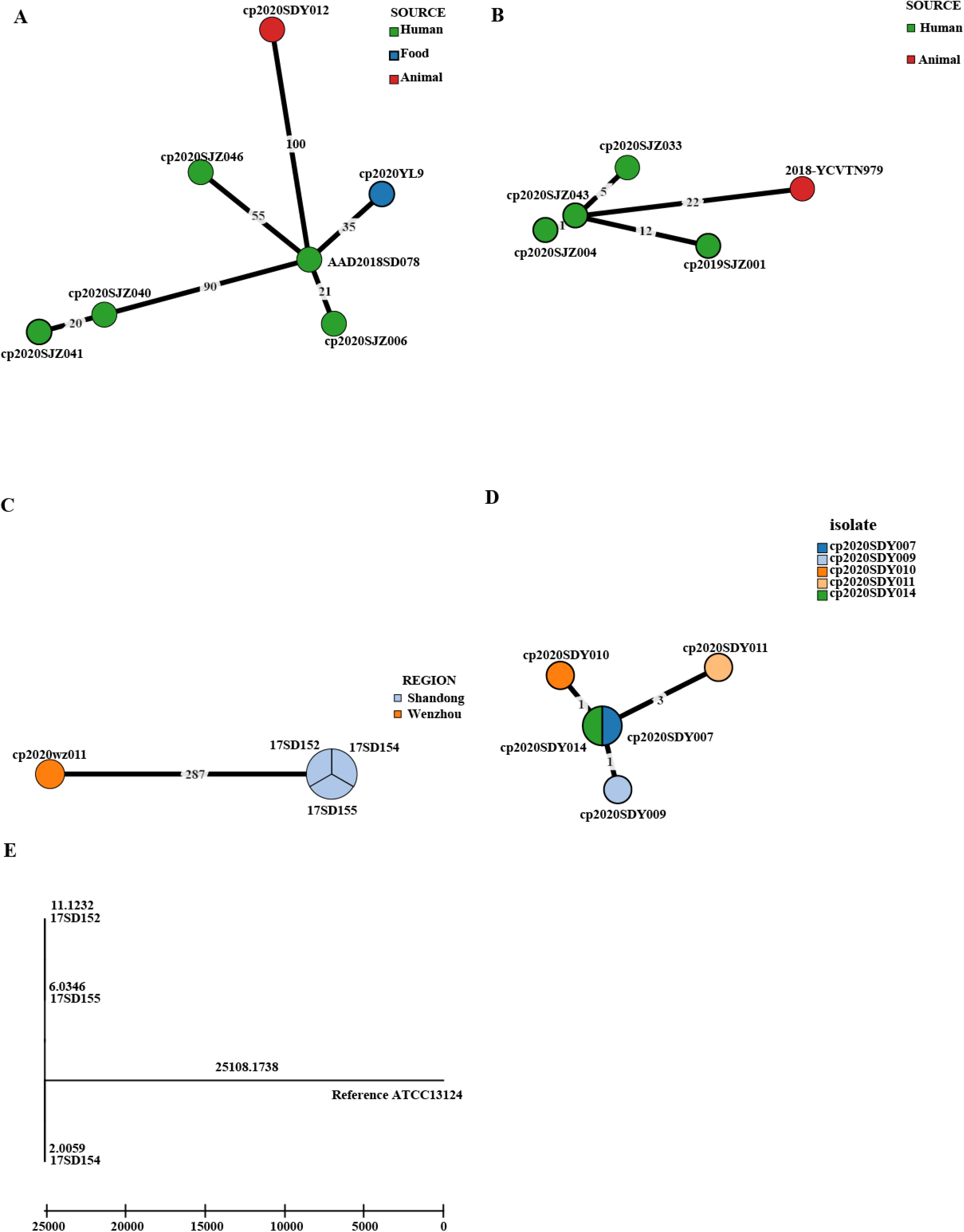
Phylogenetic tree of cgMLST and core SNP typing. (A)–(D) GrapeTree minimum-spanning tree of strains in the same STs based on the PubMLST cgMLST. The numbers on the branches represent the differences in alleles between neighboring nodes. (A) ST221, (B) ST408, (C) ST62, and (D) ST479.(E) A maximum likelihood (ML) phylogenetic tree of ST62 based on SNPs constructed by snippy and fasttree version 2.1.10 using ATCC 13124 as a reference.

In addition, within the ST408 complex, fetal cattle isolate 2018-YCVTN979 showed 22 allele differences compared to human isolate cp2020SJZ043, indicating that these isolates with identical cgMLST type were genetically heterogeneous (Fig. 4B). However, the three human clinical isolates cp2020SJZ004, cp2020SJZ043, and cp2020SJZ033 showed high genetic relatedness with ≤ 5 allele differences.

Moreover, the three AAD strains with the same ST 62 isolated from the same hospital at Shandong in 2017 (17SD152, 17SD154, and 17SD155) were also classified to the same cgMLST type showing no allele differences (Fig. 4C), strongly suggesting that they had arisen from a single source or were associated with the same transmission event. To confirm our hypothesis, cgSNP analysis was further performed for these three isolates (Fig. 4E). Within 6,914 variant sites, pairwise SNP diversity ranged from 20 to 22 SNPs for the three genomes (Supplementary Table S1), illustrating that these three isolates had clear genetic differences. Notably, there was another strain (cp2020wz011) in the ST 62 complex that displayed 287 allele differences with the above three AAD strains (Fig. 4C).

Furthermore, within the ST 479 complex, five isolates from sheep in Shandong were divided into four cgMLST types, including cp2020SDY007 and cp2020SDY014 that belonged to the same cgMLST type (Fig. 4D). The five genomes showed ≤ 3 allele differences (range 0–3) below the cutoff threshold of seven allele differences.

### Antibiotic-resistance profiles

Of the 186 strains tested, 96 were resistant to at least one antibiotic. All of the *C. perfringens* isolates showed 100% susceptibility to vancomycin and meropenem. The rates of resistance to metronidazole, penicillin, cefoxitin, moxifloxacin, and chloramphenicol were low (metronidazole: 1.08%; penicillin: 9.68%; cefoxitin: 0.54%; moxifloxacin: 6.45%; and chloramphenicol: 3.76%) (Fig. 5A). However, these isolates displayed much higher rates of antibiotic resistance to clindamycin (47.85%) (Fig. 5A).

**Figure 5.**
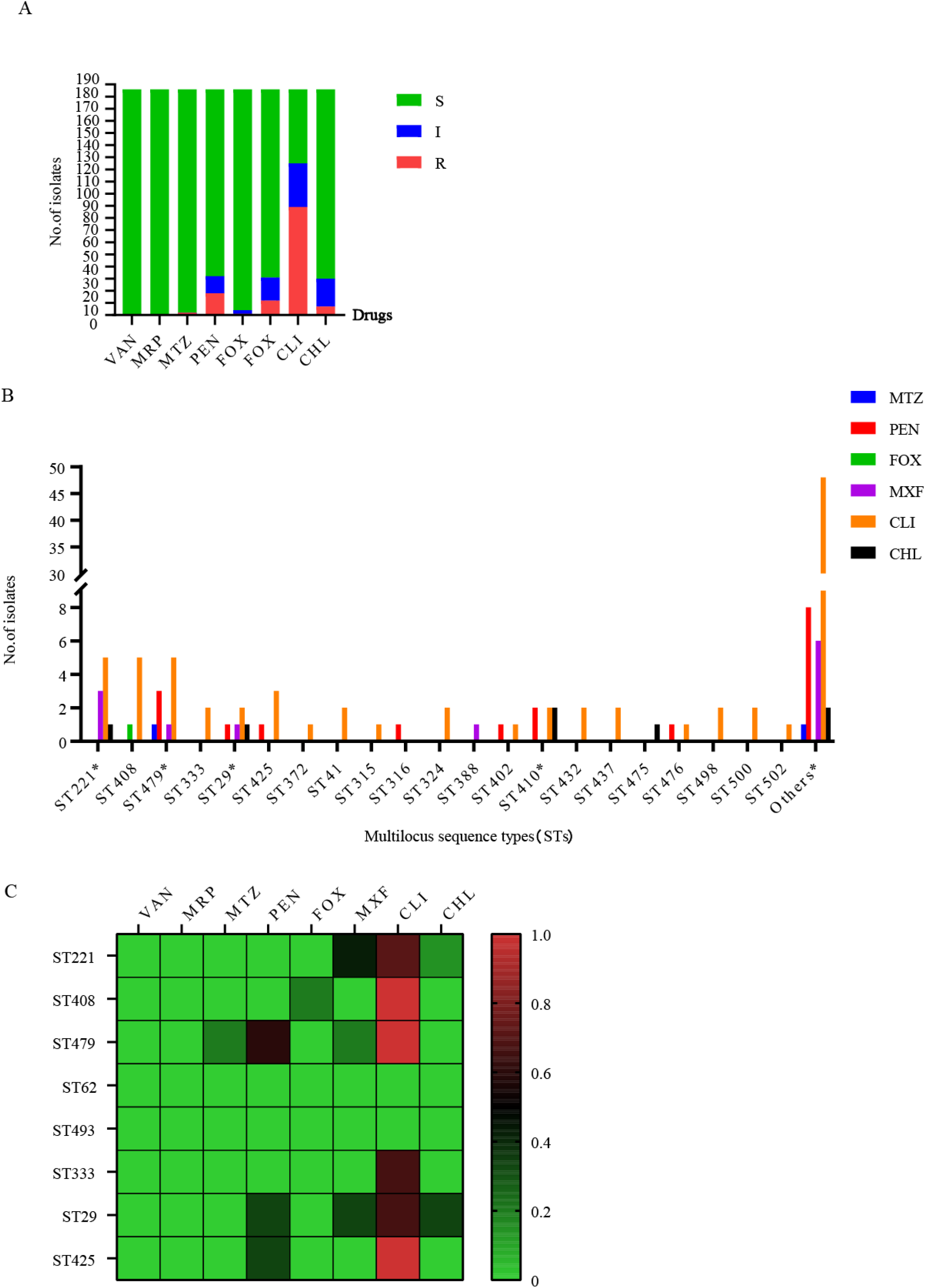
Antibiotic-resistance profiles of 186 *C. perfringens* strains. (A) Isolates sensitive, intermediate, or resistant to antimicrobial agents for 186 *C. perfringens* strains. S, sensitive; I, intermediate; and R, resistant. (B) The correlation within tested antibiotics and STs of the 186 *C. perfringens* isolates.*, MDR strain. (C) The heatmap correlating ST with antibiotic resistance. The percentages of resistance to antimicrobials are color-coded on the right side of the figure.

In the class of antibiotics (metronidazole, penicillin, cefoxitin, and moxifloxacin) with lower resistance rates, ST29 was the major molecular type of antibiotic-resistant *C. perfringens* (Fig. 5). Furthermore, there was an MDR strain (AAD2019XAzhou) in ST29 (Fig. 5B). Moreover, genotypes ST408 and ST479 were the predominant types among clindamycin-resistant isolates, followed by ST221 and ST425 (Fig. 5B). Overall, ST479, ST408, ST29, and ST410 were the most commonly identified genotypes that correlated with a high proportion of antibiotic resistance (Fig. 5C). Moreover, the strains in Group V were sensitive to all of the above antibiotics (Figs. 3 and 5B).

For isolates of human origin, all were susceptible to vancomycin, meropenem, and metronidazole. The rates of cefoxitin (0.68%), chloramphenicol (4.08%), moxifloxacin (5.44%), and penicillin (8.16%) resistance were low. The resistance rate to clindamycin (49.66%) was much higher. For isolates of animal origin, all were susceptible to vancomycin, cefoxitin, and meropenem. The antibiotics to which the tested isolates were resistant were clindamycin (52%), penicillin (24%), moxifloxacin (12%), metronidazole (8%), and chloramphenicol (4%). For isolates of food origin, all isolates were susceptible to vancomycin, meropenem, metronidazole, penicillin, cefoxitin, and chloramphenicol. The resistance rates of moxifloxacin and clindamycin were 7.14% and 21.43%, respectively (Supplementary Table S1).

The antimicrobial susceptibility testing showed that the multidrug resistance of isolates was low in this study (Fig. 5B), with a proportion of multidrug-resistant isolates of 3.23% (6/186). There were two MDR strains in ST410, one MDR strain in ST29, one MDR strain in ST221, one MDR strain in ST443, and one MDR strain in ST479 (Fig. 5B) isolated from sheep (n = 1) and humans (n = 5). The MDR proportion of human isolates was 3.40% (5/147). The MDR proportion of animal isolates was 4% (1/25). There were no multidrug-resistant strains of food origin in our study. Supplementary Table S1 contains a complete list of antibiotic-resistance profiles found in the study isolates.

## Discussion

To the best of our knowledge, this is the first relatively comprehensive cross-sectional study concerning the molecular characterization of *C. perfringens* isolates in China. In our study, samples from patients, healthy humans, animals, and food were subjected to MLST analysis, toxin type analysis, cgMLST analysis, and antibiotic susceptibility testing.

The MLST analysis of Chinese strains indicated a wide genetic diversity of *C. perfringens*, as previously reported (17). In our study, the predominant STs from human isolates (ST221, ST62, ST408) were significantly distinct from the predominant STs (ST479) from animal isolates, a result that was consistent with previous research on global *C. perfringens* strains, indicating a diverse distribution of *C. perfringens* among multiple sources (18, 19). In global surveys, a clonal complex (CC) represented by three STs (ST 98, ST 41, and ST 110) predominantly representing type F (18/20 strains) was mostly associated with human illnesses (18). Among the predominant STs, ST 54 was associated with enteritis cases in foals and dogs, and ST 58 was associated with necrotic enteritis in poultry (18). In our study, most type F strains (10/11 strains) were associated with human illnesses, different from the results of previous studies.

In terms of strains of human origin, the isolates recovered from human diarrhea cases (n = 111) were found to be of type A (n = 102), type F (n = 8), and type D (n = 1). Type A and type F were the major toxin types in the isolates, basically in accordance with earlier reports (20, 19). Interestingly, there was one isolate of type D that carried *cpe* and/or *cpb2* genes, indicating that it had the potential to produce the CPE toxin (1). ST221 was the dominant genotype of *C. perfringens* among the isolates, with significant genetic diversity. Previous results revealed that ST41 and ST77 were associated with human illnesses (18, 19). Consistent with these findings, we observed that the strains ST41 (2/2, type A) and ST77 (2/2, type F) were isolated from patients with diarrhea. The resistance rates of *C. perfringens* depend on the use of antibiotics in different regions. For example, some studies in Hungary, Slovenia, and northern Taiwan showed that cefoxitin, meropenem, and metronidazole were successful against *C. perfringens*. Only a minority were penicillin (2.6%) or clindamycin (3.8%) resistant (21). However, in our study, we found high resistance (49.66%) to clindamycin in human origin samples in China. Similarly penicillin-resistant (10.2%) and penicillin-insensitive strains (18.2%) were much more frequent in this study than in previous studies. Moreover, recent data from Italy showed that clindamycin resistance was >20% for *Clostridium* spp. (22). In fact, penicillin and clindamycin have long been considered excellent drugs for *Clostridial* infections. In particular, penicillin is currently the drug of first choice for *C. perfringens* infections. Thus, our findings suggest that the two antibiotics should be used with caution as empirical therapies in China.

Among the strains of animal origin, all were found to be type A in this study. A relatively high prevalence of MDR (73.4%) was found among the isolates from different animals in a previous study (24). The MDR profile was resistant against tetracycline (84%), erythromycin (72%), sulfamethoxazole (97.3%), trimethoprim (96.3%), and streptomycin (84.4%) (24). Interestingly, the prevalence of MDR *C. perfringens* strains was rather low in this study (4%, 1/25 animals), possibly due to different choices of tested antimicrobials. The small scale of animal samples or low use of antibiotics in selected farms is also potential cause. Furthermore, the environment in which *C. perfringens* lives (e.g., the gut) may play a significant role in drug resistance. Therefore, in the future, close monitoring is necessary for a more thorough investigation of antibiotic resistance of *C. perfringens* in China. Rational antibiotic use may play an important role in reducing the drug resistance of *C. perfringens*.

*C. perfringens* is an important microorganism as a foodborne pathogen. Among the *C. perfringens* of food origin, in a previous study in Egypt, most of the food origin isolates (74%) (beef, chicken meat, and raw milk) exhibited MDR patterns. Strains isolated from raw camel milk in Isiolo County, Kenya, had high resistance to chloramphenicol (42.37%), kanamycin (40.68%), tetracycline (37.29%), and gentamycin (35.59%) (25). Another study in Korea reported that resistance to chloramphenicol (26/38, 68.4%) and metronidazole (13/38, 34.2%) was observed in *C. perfringens* from retail meats (26). Surprisingly, in our study, all of the isolates from retail meat products were susceptible to chloramphenicol and metronidazole, and none of the isolates showed an MDR pattern. Given that most of the samples in this study were of human origin, the selected antimicrobials were biased toward clinical use, a feature that was inconsistent with other studies using isolates of animal or food origin.

The three isolates (17SD152, 17SD154, and 17SD155) with the same ST62 isolated from the same hospital were assigned to the same cgMLST type (with no allele difference). Their visit times were close, no more than half a month apart. The three patients were 6 months old, 34 years old, and 62 years old. The three isolates collected at a single Shandong hospital inferred a potential transmission chain. The five isolates within the ST 479 complex (cp2020SDY007, cp2020SDY009, cp2020SDY010, cp2020SDY011, and cp2020SDY014) of sheep origin in Shandong were divided into four cgMLST types with allele differences range 0–3, including cp2020SDY007 and cp2020SDY014 that belonged to the same cgMLST type (Fig. 4D). Thus, we strongly suspect that transmission of these *C. perfringens* strains in single farm took place. Genetic relatedness of strains isolated from three AAD patients from the same hospital and five sheep from the same farm in Shandong indicates that these isolates are originated from a single source or associated with the same transmission event and inter-host transfer of *C. perfringens*.

In our study, sequence type complex 221 was further divided into different cgMLST profiles. The STc221 strains across different provinces and hosts showed large allelic differences, indicating independent development and distribution. Meanwhile, this pattern showed that cgMLST had higher discrimination than the classical MLST method. Overall, cgMLST made it easy to clarify transmission routes, target the epidemiological investigation, and delineate infection control interventions. The cgSNP based on whole-genome sequencing (WGS) has advantages in understanding the dissemination of *C. perfringens*.

In conclusion, this study provides a glance at the molecular features and genetic characters of *C. perfringens* isolates in China. Using MLST analysis of 186 *C. perfringens* strains from diverse hosts, we found distinct differences in dominant STs among humans and animals. Similar characters were reported in other studies (26). Moreover, we gained insight into the antibiotic-resistance profiles in China of this significant pathogen. In our study, the percentage of MDR strains was much lower than previously reported. Interestingly, isolates of human origin were highly clindamycin-resistant and penicillin-insensitive, implying that their use as empirical therapies in China should be approached with caution. We analyzed the genetic relatedness of these specific complexes with the same STs to identify potential transmission links based on MLST, cgMLST, and cgSNP. Our results indicated that the isolates of the same ST may be genetically unrelated. cgMLST and cgSNP are more discriminant than traditional MLST and are suitable approaches for outbreak and transmission analyses. Further studies focused on *C. perfringens* of animal and food origin nationwide are needed to gain deep insights into prevention and control strategies, routine outbreak detection, and national surveillance in China.

## Materials and methods

### Sample collection

A total of 186 *C. perfringens* strains were recovered and cultivated in our lab and characterized using the following molecular techniques. These isolates were obtained from people, including patients and healthy people, domestic animals (sheep, calves, and pigs), and food from nine different regions across China, from 2013 to 2021 (Fig. 2). There were 47 isolates from Shijiazhuang, 81 isolates from Shandong, 16 isolates from Yunnan, three isolates from Sichuan, 11 isolates from Wenzhou, 14 isolates from Yulin, six isolates from Xi’an, two isolates from Hunan, and six isolates from Beijing (Fig. 2). Among these 186 strains, 139 strains were from clinical sources; 24 strains were from animal sources; and 14 were from food sources, comprising 174 type A strains (93.55%), 11 type F strains (5.91%), and one type D strain (0.54%) (Fig. 2).

### Isolation and identification of *C. perfringens*

After ethanol shock treatment, all fecal specimens were inoculated with 5% egg yolk on selective tryptose sulfite cycloserine (TSC) agar (Oxoid, Basingstoke, UK) plates and incubated at 37°C for 48 hours in an anaerobic jar (Mart, NL). Following vacuuming, an anaerobic atmosphere of 80% nitrogen, 10% hydrogen, and 10% carbon dioxide was injected. *C. perfringens* colonies were identified based on biochemical tests, colony and cell morphology, and other characteristics, and the 16S rRNA gene was used to identify the bacteria (short Gram-positive bacilli, dual hemolysis, and gelatinase and pectinase-producing bacteria) (27).

### DNA extraction

All isolates used in this study were maintained on BHI plates with 5% sheep agar under anaerobic conditions. The genomic DNA of these isolates was extracted using a bacterial genomic DNA extraction kit (Qiagen, Germany) according to the manufacturer’s instructions. DNA was then dissolved in 100 μl of TE and stored at −20° C until use.

### MLST and phylogenetic analysis

The eight gene loci (*colA, groEL, sodA, plc, gyrB, sigK, pgk, and nadA*) listed in the public *C. perfringens* MLST database (https://pubmlst.org/organisms/clostridium-perfringens) were used in this study (27). PCR assays were performed in final volumes of 25 μl containing 12.5 μl of 2 × PCR Taq Mastermix (MgCl, dNTP, Taq enzyme) (Takara Bio Inc., Japan); 0.5μl of each primer (10 mmol/L); 1.5 μl of DNA template; and double-distilled water for a final volume of 25 μl. Reactions were performed with initial denaturation at 94° C for 2 min, followed by 35 cycles at 94° C for 30 s, at 55° C for 60 s and at 72° C for 60 s, and a final extension at 72° C for 8 min. Then, the PCR products were purified using a commercial kit (Omega Bio-Tek, USA) and were sent for Sanger nucleotide sequencing in both directions (Beijing DIA-UP BIOTECH Co., Ltd., China).

To determine the sequence type (ST) of all the tested 186 strains of *C. perfringens*, DNA sequences were submitted to a public *C. perfringens* MLST database (https://pubmlst.org/organisms/clostridium-perfringens) (29). New alleles and STs were assigned in the *C. perfringens* MLST database. The minimum-spanning tree was constructed using BioNumerics software version 5.1.

### cgMLST and evolutionary relationship analysis

A *C. perfringens* cgMLST scheme comprising 1,431 target loci based on genomic sequences was used, and neighbor-joining was used to create dendrograms based on cgMLST information via the PubMLST website (https://pubmlst.org/organisms/clostridium-perfringens).

An alignment of SNPs in the core genome using ATCC13124 as a reference was constructed using Snippy (version 4.6). The size of the core genome for *C. perfringens* isolates included in this study was 22,493 bp, and the average size of the *C. perfringens* genome for isolates sequenced in this study was 3.26 Mb. A phylogenetic tree was produced using the 3,256,683 bp core genome SNP alignment, and the maximum likelihood tree was constructed using fasttree (version 2.1.10).

### Antimicrobial susceptibility test

The E-test method was applied for antimicrobial susceptibility analysis (Liofilchem s. r. l., Roseto degli Abruzzi, Italy) of the *C. perfringens* isolates. The following eight antimicrobials were tested: clindamycin, chloramphenicol, vancomycin, cefoxitin, meropenem, metronidazole, penicillin, and moxifloxacin. Isolates were suspended in Brucella broth to achieve a turbidity of 1.0 McFarland, and then inoculated onto Brucella agar supplemented with 5% sheep blood, 1 mg/L Vitamin K1, and 5 mg/L Hemin Chloride. Plates were incubated at 37° C for 18–24 h in anaerobic conditions. *C. perfringens* ATCC 13124 was used as a quality control. MDR was characterized as resistance to at least one agent in three or more antimicrobial classes (30).

The interpretation criteria for Gram-positive anaerobic bacteria are available from the Clinical and Laboratory Standards Institute (CLSI) (31) and the European Committee on Antimicrobial Susceptibility Testing (EUCAST) (32). MIC breakpoints for clindamycin (≥ 8 μg/ml), chloramphenicol (≥ 32 μg/ml), cefoxitin (≥ 64 μg/ml), meropenem (≥ 16 μg/ml), metronidazole (≥ 32 μg/ml), penicillin (≥ 2 μg/ml), moxifloxacin (≥ 8 μg/ml) (31), and vancomycin (> 2 μg/ml) (31) were used to read the breakpoints.

## Supplementary data

SUPPLEMENTAL FILE 1, XLSX file, 33 KB.

SUPPLEMENTAL FILE 2, XLSX file, 14 KB.

## Acknowledgments

We thank Haijian Zhou, Yutong Kang, Jie Che, and Zhizhou Tan for their suggestions in both experiments and manuscript preparation.

This work was supported by the National Key Research and Development Program of China (2021YFC2301000).

We thank LetPub (www.letpub.com) for its linguistic assistance during the preparation of this manuscript.

